# Magnetic Enrichment of Immuno-Specific Extracellular Vesicles for Mass Spectrometry Using Biofilm-Derived Iron Oxide Nanowires

**DOI:** 10.1101/2022.05.01.490183

**Authors:** Quang Nghia Pham, Marnie Winter, Valentina Milanova, Clifford Young, Mark R. Condina, Peter Hoffmann, Nguyen T. H. Pham, Tran Thanh Tung, Dusan Losic, Benjamin Thierry

## Abstract

Immuno-specific enrichment of extracellular vesicles (EVs) can provide important information into cellular pathways underpinning various pathologies and for non-invasive diagnostics, including mass spectrometry-based analyses. Herein, we report an optimised protocol for immuno-magnetic enrichment of specific EV subtypes and their subsequent processing with liquid chromatography-tandem mass spectrometry (LC-MS/MS). Specifically, we conjugated placental alkaline phosphatase (PLAP) antibodies to magnetic iron oxide nanowires (NWs) derived from bacterial biofilms and demonstrated the utility of this approach by enriching placental specific EVs (containing PLAP) from cell culture media. We demonstrate efficient PLAP+ve EV enrichment for both NW-PLAP and Dynabeads™-PLAP, with high PLAP protein recovery (83.7±8.9% and 83.2±5.9%, respectively), high particle-to-protein ratio (7.5±0.7×10^9^ and 7.1±1.2×10^9^, respectively), and low non-specific binding of non-target EVs (7±3.2% and 5.4±2.2%, respectively). Furthermore, our optimized EV enrichment and processing approach identified 2518 and 2545 protein groups with LC-MS/MS for NW-PLAP and Dynabead™-PLAP, respectively, with excellent reproducibility (Pearson correlation 0.986 and 0.988). These findings demonstrate that naturally occurring iron oxide NWs have comparable performance to current gold standard immune-magnetic beads. The optimized immunospecific EV enrichment for LC-MS/MS method provides a low-cost and highly-scalable yet efficient, high-throughput approach for quality EV proteomic studies.

## 1. Introduction

Extracellular vesicles (EVs) are a collective group of lipid bilayer-enclosed particles present in most biological fluids such as blood, amniotic fluid, urine, cerebrospinal fluid and cell culture media.^[1–3]^ EVs are important in many biological processes including intercellular communication, signal transduction, immune response and maintenance of cellular homeostasis.^[4–5]^ EVs have been increasingly recognized as a promising source of biomarkers since the biomolecule cargo including DNA, mRNA, proteins, lipids and microRNA, often reflect the physiological and pathological conditions of the parent cells.^[6]^ Moreover, biomolecules are better protected from enzymatic degradation while being transported within EVs as compared to free proteins and RNAs.^[7–9]^ Given that EVs can be non-invasively obtained through blood serum and plasma, analyses of EV number and molecular content have already provided important insights for the understanding and diagnosis of diseases.^[10–14]^

According to the Minimal Information for Studies of Extracellular Vesicles 2018 (MISEV2018), EVs can be classified as small (< 200 nm) and large EVs (> 200 nm).^[15]^ Ultracentrifugation and size exclusion chromatography are the most popular approaches used to enrich EVs based on their physicochemical properties, namely density and size.^[16]^ Importantly, EVs display specific proteins which relate to their biogenesis and cell of origin. These proteins can be exploited for immunospecific enrichment of EV subtypes.^[17–18]^ To this end, affinity ligands such as antibodies and aptamers are typically immobilized on a solid support allowing for selective enrichment.^[19]^ The most common immunoaffinity approach is based on magnetic materials conjugated with antibodies to bind specific EV proteins followed by separation under a magnetic field. Immuno-magnetic methods produce high-purity isolates of immuno-specific EVs with low contamination rates in a rapid and efficient manner.^[20–23]^ Immunoaffinity EV enrichment can also be achieved with microfluidic devices.^[24]^ However, microfluidic approaches typically suffer from low throughput which limits their use for applications that require a large amount of EVs for downstream analyses including biomarker discovery, particularly when low-abundant EV subtypes are the target.^[25–26]^ Immuno-magnetic isolation therefore currently offers the best approach for the enrichment of EV subtypes and has been shown to be compatible with proteomic approaches such as liquid chromatography-tandem mass spectrometry analysis (LC-MS/MS).^[27]^

Many types of magnetic materials have been investigated for immuno-magnetic EV enrichment. These materials are typically selected based on their responsiveness to magnetic fields and high surface area for binding. Iron oxide-based particles are the most common and a range of sizes have been used including micron sized magnetic beads^[25]^ (e.g. Dynabeads™)^[28]^ and nanoparticles.^[29]^ The availability of commercially available beads such as Dynabeads™ and automated technologies has made their use more widespread and standardized, with well-established protocols available for many applications.^[34]^

Importantly, a necessary trade-off should be established with respect to the dimension of the magnetic materials used for EV enrichment. On one hand, high surface area such as in the case of magnetic nanoparticles enables high EV capture yield. On the other hand, larger materials such as in the case of magnetic microbeads afford better magnetic recovery of the EVs and their biological cargo. Magnetic nanowires (NWs) offer an excellent compromise balancing the requirements of high surface area (for efficient capture of target EVs) and strong magnetic strength (for efficient magnetic recovery).^[35]^ Several magnetic NWs have shown utility in a range of bioanalytical applications, including the enrichment of cells, proteins, RNA, DNA and EVs.^[30, 33, 36]^ For example, magnetic poly pyrrole NWs (electrochemically prepared from an anodized aluminum oxide (AAO) template by embedding iron oxide magnetic nanoparticles) allowed rapid enrichment of EVs with relatively high yield and purity when conjugated to a cocktail of EV specific antibodies (anti-CD9, anti-CD63, and anti-CD81), with a threefold increase in the yield compared to commercially available methods such as ExoQuick™ and Invitrogen Total Exosome Isolation Kit.^[31]^ The same approach could also be used to achieve rapid and high yield capture of cell-free DNA and circulating tumor cells while minimizing DNA loss and damage.^[37]^ The *in vitro* cellular uptake and internalization of magnetic nickel NWs has also been shown to be a fast and cost-effective approach for non-specific EV enrichment from cell culture.^[33]^ While these studies convincingly outline the potential of magnetic NWs, the reliance on complex, expansive and inherently low-throughput synthesis routes such as templating hampers the routine use of magnetic NWs for EV enrichment. A simple, cost-effective and scalable method to prepare magnetic NWs for immunomagnetic enrichment of EVs and other relevant biomolecules is extremely desirable.

Towards addressing this challenge, we report here an immuno-magnetic enrichment method for specific EV subtypes and their subsequent analysis with mass spectrometry. We benchmarked the enrichment performance of naturally produced iron oxide NWs with commercially available Dynabeads™ and perform mass spectrometry on enriched EVs. The magnetic NWs are extracted and purified from biofilms of zetaproteobacteria *Mariprofundus ferrooxydans* through an environmentally friendly process.^[38]^ These NWs have previously been shown to have a large surface to volume ratio, respond rapidly to magnetic fields, and have been explored as magnetically targeted drug carriers.^[38]^ For bioconjugation with antibodies and improved colloidal stability, we modified the NWs with reversible addition-fragmentation chain transfer (RAFT)s diblock copolymers (*p*AAm) with high binding affinities to iron oxide (**Figure 1A**). As a proof of concept, we demonstrated the immuno-specific enrichment of EVs based on the expression of placental alkaline phosphatase (PLAP) (Figure 1B). PLAP is expressed in the human placenta mainly by syncytiotrophoblasts and is known to be present in maternal blood with levels increasing over gestation.^[39–40]^ Syncytiotrophoblasts derived EVs are anticipated to provide a non-invasive window into placental (and hence fetal) health.^[41]^ However, compared to maternal EVs, PLAP+ve EVs represent only a portion of total EVs in the maternal circulation (up to ~20% in early pregnancy),^[42]^ meaning that efficient immuno-enrichment is required in order to fully understand the relevance of these vesicles or harness their diagnostic utility. The magnetic NWs conjugated with anti-PLAP antibodies efficiently enriched PLAP+ve EVs produced by the trophoblast-like chorionic carcinoma BeWo cell line used as a model, similar to that afforded by the current commercial gold standard Dynabeads™. In addition, we also designed and optimized a protocol for LC-MS/MS analysis of the enriched PLAP+ve EVs.

**Figure 1.**
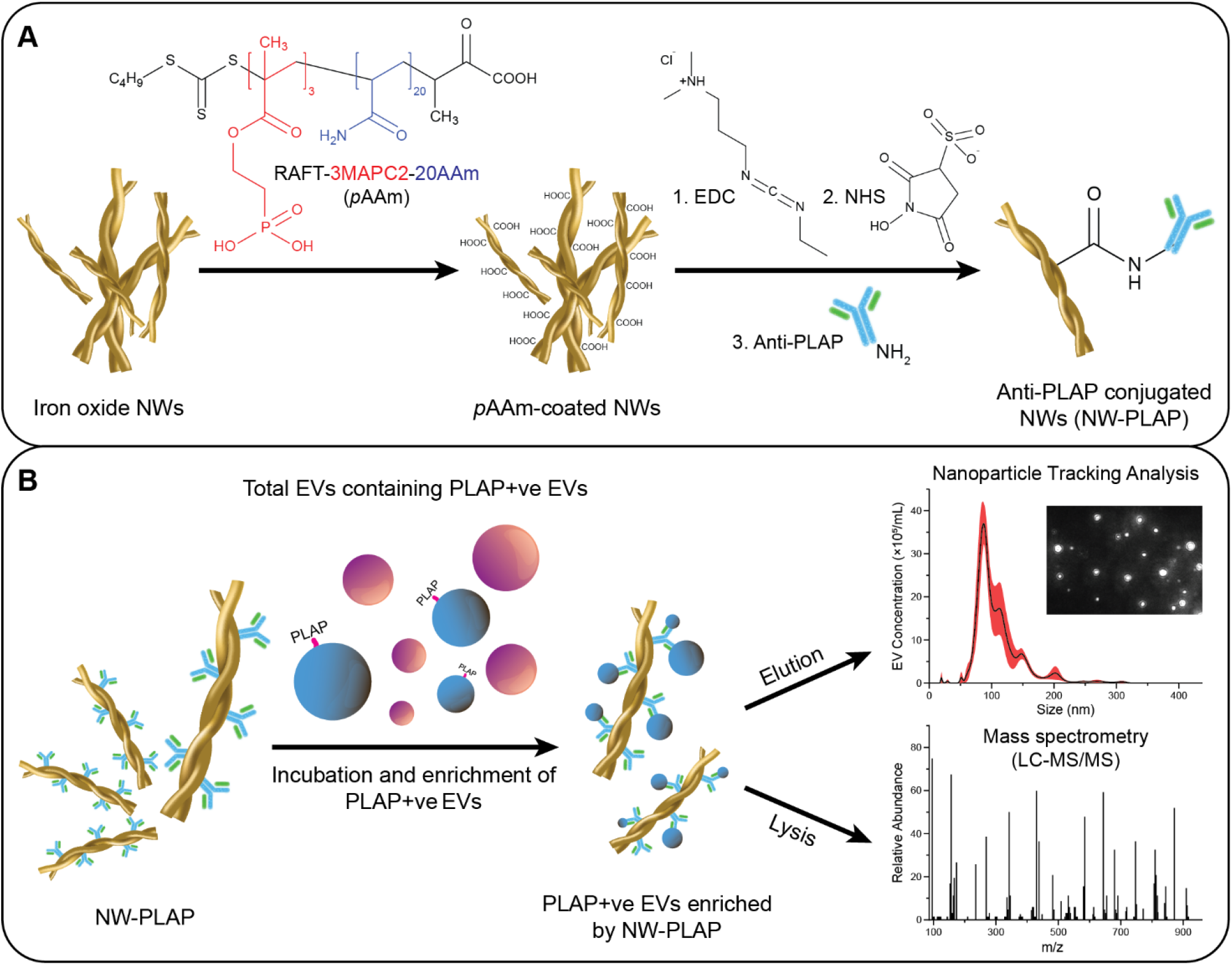
Surface modification of iron oxide NWs and enrichment of PLAP+ve EVs. (A) Schematic of biofunctionalization route of the NWs with (RAFT)s diblock copolymers (poly(methacryloyloxyethyl phosphate)3-block-poly(acrylamide)20 RAFT polymer, referred to as *p*AAm) and anti-PLAP antibodies. (B) Enrichment procedure of PLAP+ve EVs for downstream analyses with nanoparticle tracking analysis and mass spectrometry (LC-MS/MS).

## 2. Results and Discussion

### 2.1. Characterisation of iron oxide NWs

The morphologies of the pristine iron oxide NWs and NW-*p*AAm are presented in **Figure 2**. The scanning electron microscope (SEM) analysis revealed that the bacteria-derived NWs used in this study appeared as helical-like structures comprising of twisted NWs (Figure 2A). These NWs varied in size, ranging approximately 460–950 nm with a polydispersity of 0.62 as estimated by dynamic light scattering (DLS) (Figure S1A). The average diameter of a single NW was 66.7±16.7 nm as measured by SEM. The transmission electron microscope (TEM) image (Figure S1B) showed the dimensions and morphology of iron oxide NWs which are consistent with SEM images (50–80 nm). Their polycrystalline structures with lattice refractions are corresponding to a mixture of magnetite and hematite (Figure S1C). The magnetic property studied by the measurements of magnetic hysteresis revealed that the iron oxide NWs exhibited a ferromagnetic behavior (Figure S1D). NWs coated with the diblock copolymers (NW-*p*AAm) showed similar wire-like structures (Figure 2B) with increased size distribution as seen by the increased scattering in the DLS measurements (688±22 nm for NWs and 1110±26 nm for NW-*p*AAm) (Figure S1A). Note that diameters from the DLS measurements should only be considered coarse approximations considering the non-spherical nature of the NWs.^[43]^ After the conjugation of antibodies, the approximated hydrodynamic size of NW-IgG increased to 1154±19.5 nm with a polydispersity of 0.44 (Figure S1A). Dynabeads were also imaged with SEM to confirm their homogeneous spherical morphologies. A mean size of 2.7 μm and a polydispersity of 0.093 were measured by DLS (Figure S2).

**Figure 2.**
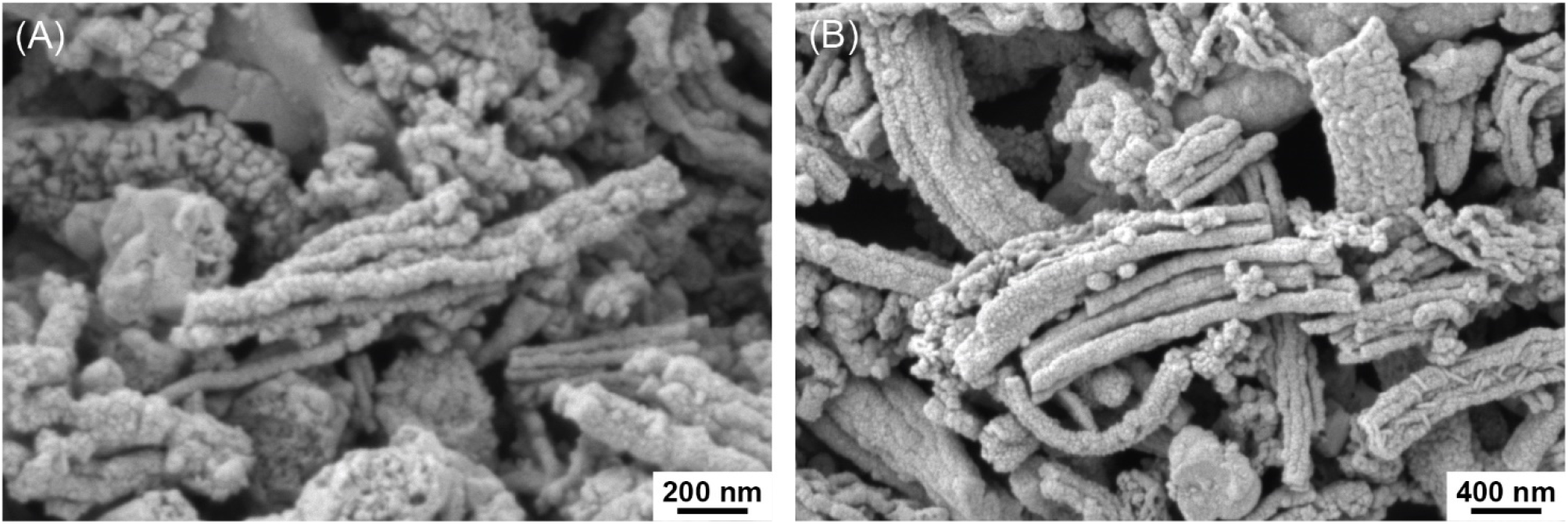
SEM images of (A) NW and (B) NW-*p*AAm.

To further confirm the successful functionalization of the NWs with carboxylic acid groups and conjugation with antibodies, chemical analyses with X-ray photoelectron spectroscopy (XPS) were performed for NWs, NW-*p*AAm and NW-IgG (**Figure 3**). As shown in Figure 3A, elemental survey scans contained peaks attributed to Fe 2p at 711.7 eV (Fe 2p3/2) and 725.1 eV (Fe 2p1/2) in all three samples, which are the characteristics of Fe (III) of α-Fe2O3.^[44–45]^ A peak at the lower binding energy of approximately 708 eV corresponding to Fe (II) was not observed. These features were in accordance with previous results conducted by X-ray diffraction (XRD) and Fourier-transform infrared (FTIR) analyses of iron oxide NW-derived from *M. ferrooxydans^[46]^*. Quantification of the elemental composition of the three samples (**Table 1**) showed increases in the intensities of carbon (C 1s at 284.8 eV), oxygen (O 1s at 530 eV), and nitrogen (N 1s at 399.4 eV), which indicate successful coating of the NWs with the copolymers and further conjugation with antibodies. Specifically, compared to the NWs, both NW-*p*AAm and NW-IgG showed significant increases in elemental carbon contents (39.51% for NW-*p*AAm and 47.79% for NW-IgG vs. 30.67% for NW), which was mainly attributed to increased hydrocarbon indicated by the CC/CH component at 284.8 eV in the high-resolution scans (Figure 3B). The increases in the intensity of CN (285.6 eV) and changes in the intensity of the CO (286.3 eV), C=O (288.1 eV), and O-C=O (289 eV) components can also be attributed to the presence of the copolymer on the surface of the NWs. Deconvolution of the O 1s high-resolution spectra in Figure 3C shows the presence of the expected Fe-O component at approximately 529.7 eV in all three samples. After coating and antibody conjugation, the intensities of the O=C component (531.6 eV) increased, which is in agreement with the changes observed in the C 1s spectra. While nitrogen was not detected in the pristine NWs (Figure 3D), N 1s high-resolution spectra showed a prominent peak attributed to CN at approximately 399.8 eV for both NW-*p*AAm and NW-IgG. As expected, the N% substantially increased after conjugation with antibodies (3.34% for NW vs. 8.13% for NW-IgG). Using 1 mg of NWs and 40 μg of antibodies, the conjugation efficiency was determined to be 89%. As the NWs used in this study are heterogeneous in size and shape, the batch-to-batch variation of the surface modification and antibody conjugation was also examined. As shown in Table S1, minimal variations in the XPS elemental composition percentages were observed after *p*AAm coating and antibody conjugation for two independent NW batches, demonstrating the reproducibility of the modification and conjugation processes.

**Figure 3.**
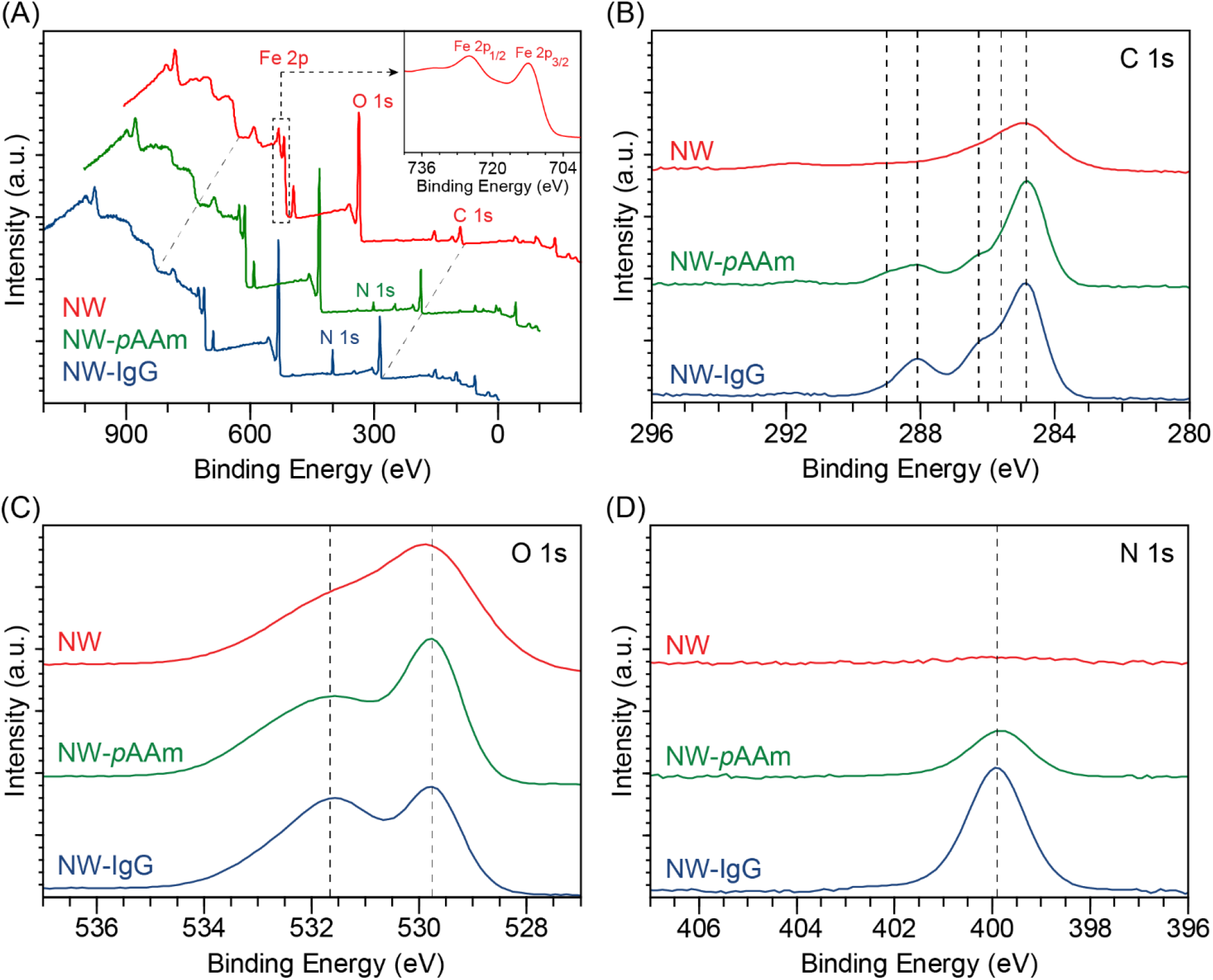
XPS analyses of NW, NW-*p*AAm, and NW-IgG. (A) Survey scan spectra with inset displaying Fe 2p peak. (B) High-resolution spectra of C 1s. Dashed lines indicate major peaks of CC/CH (284.8 eV), CN (285.6 eV), CO (286.3 eV), C=O (288.1 eV), and O-C=O (289 eV). (C) High-resolution spectra of O 1s. Dashed lines indicate major peaks attributed to Fe-O (529.7 eV) and O=C (531.6 eV). (D) High-resolution spectra of N 1s. Dashed line indicates CN (399.8 eV).

**Table 1.** Summary of XPS elemental and chemical composition.

| NW samples | XPS elemental composition (%) |  |  | C 1s components (%) |  |  |  |  | O 1s components (%) |  |
| --- | --- | --- | --- | --- | --- | --- | --- | --- | --- | --- |
|  | C | N | O | CC/CH | CN | CO | C=O | O-C=O | Fe-O | O=C |
| NW | 30.67 | - | 69.33 | 53.82 | - | 28.94 | 10.56 | 6.68 | 56.49 | 43.51 |
| NW-pAAM | 39.51 | 3.34 | 57.15 | 64.26 | 3.83 | 17.04 | 11.25 | 3.62 | 34.21 | 65.79 |
| NW-IgG | 47.79 | 8.13 | 44.09 | 61.12 | 9.8 | 11.34 | 15.34 | 2.4 | 25.78 | 72.42 |

“-” indicates below the detection limit

### 2.2. PLAP+ve EV enrichment performance

To assess the EV capture performance of the iron oxide NWs and Dynabeads, EVs enriched from the culture media of the BeWo cell line were used as model. BeWo EVs contain PLAP on their lipid bilayer membranes, with PLAP+ve EVs accounting for approximately 28.8% of the total EVs as quantified by fluorescent nanoparticle tracking analysis (NTA),^[47]^ making BeWo a good model for testing the enrichment of immuno-specific EVs. Due to the comparatively low number of EVs in cell culture media, an ultracentrifugation step was first used to concentrate total BeWo EVs (defined as the “pre-enriched” sample). Specific amounts of NW-PLAP and Dynabead-PLAP were then added to 250 μL of the pre-enriched BeWo EVs containing 44.5 μg mL^-1^ of proteins. The magnetic NWs are readily collected using an external magnetic field, in agreement with a previous report.^[38]^

We used NTA to characterize the size distribution and concentration of the PLAP+ve EVs enriched by both NW-PLAP and Dynabead-PLAP. Specifically, EVs were eluted by Glycine-HCl prior to NTA analysis as it is not possible to enumerate bound EVs with NTA. For all other EV analysis methods reported, EVs were lysed directly after immuno-magnetic enrichment without elution. Although elution efficiency was not specifically examined, the size distribution of the eluted EVs was 62–245 nm with the mean/mode size of 147.2±2.6/108.7±14.5 nm and 132.4±10.6/113.8±13.2 nm in the case of the NWs and Dynabeads *(p* > 0.05), respectively (Figure S3A-C). This data indicates that enriched PLAP+ve EVs include both small and large EVs.

We then further investigated the EV capture efficiency for the magnetic NWs by varying the amount of NW-PLAP used (0.02, 0.05, 0.1, 1 and 2 mg) and the enrichment time (10, 30, 60, 120 and 240 min). EVs captured on the NW-PLAP were lysed and the amount of PLAP protein in the lysate was referenced to the amount in the pre-enriched EVs solution. As the amount of NW-PLAP increased from 0.02 to 1 mg, the PLAP recovery plateaued at approximately 80% from 0.5 mg of NW-PLAP (**Figure 4**A). We therefore selected 0.5 mg of NW-PLAP per sample for subsequent optimization experiments. As shown in Figure 4B, the recovery of PLAP increased from 35% to 82% as the time increased from 10 min to 240 min. The optimal enrichment time was 60 min (83.7±5.7%), with prolonged incubations leading to no further improvement in PLAP recovery. The optimized sample processing method was therefore, selected to include a 60 min incubation for the recovery of PLAP+ve EVs in a 250-μL sample (corresponding to 44.5 μg mL^-1^ of proteins) with 0.5 mg of NW-PLAP.

**Figure 4.**
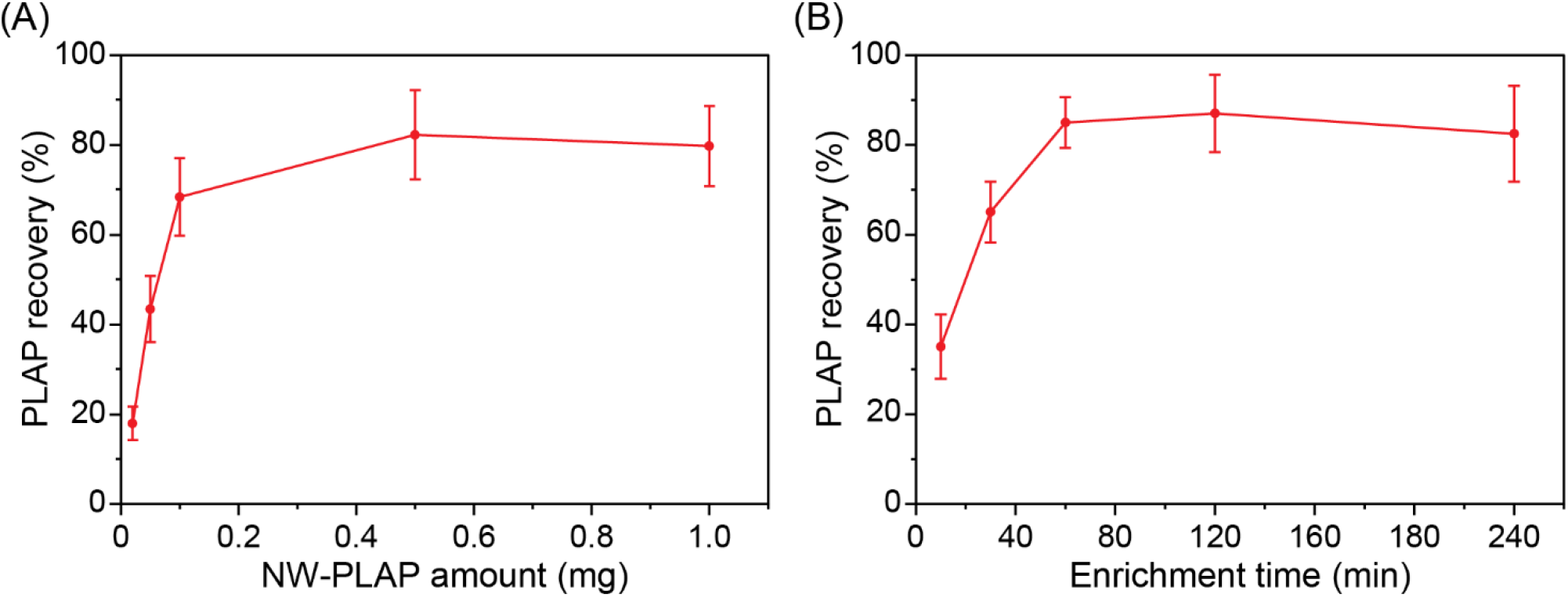
Effect of (A) the amount of NW-PLAP (0.02, 0.05, 0.1, 0.5 and 1 mg) and (B) enrichment time (10, 30, 60, 120 and 240 min) on the recovery of PLAP proteins from model BeWo EV samples. Data are presented as means ± standard deviation (SD) (*n* = 3).

Having determined the optimum recovery conditions for the anti-PLAP antibody-conjugated magnetic NWs, we then benchmarked their performance against the broadly used commercially available Dynabeads. Performance comparisons were performed using the same weight of NWs and Dynabeads™ (1 mg). The amount of soluble protein co-enriched with EVs was first determined by measuring the particle to protein ratios described previously as a measure of EV purity.^[48]^ Using NTA to quantify the number of eluted EVs after immuno-magnetic enrichment and bicinchoninic acid (BCA) assay to quantify the total protein concentrations in the lysates, the particle to protein ratio for the NWs (2.84±1.12×10^10^) was found to be comparable to that of the Dynabeads (2.53±1.1×10^10^) *(p* > 0.05) (Figure S3D). Both ratios were in the range expected of a pure EV preparation (> 2×10^10^)^[49]^ and comparable to purity ratios previously reported for immuno-magnetic EV enrichment from cell culture media^[25]^ and plasma,^[50]^ confirming that both magnetic NWs and Dynabeads can be used to enrich EVs with high purity.

We then evaluated the specificity of the NW-PLAP and Dynabead-PLAP in terms of non-specific EV-binding. PLAP+ve EVs were recovered from the pre-enriched total BeWo EVs (prepared as described above) using the same weight of NWs and Dynabeads (1 mg) conjugated with either anti-PLAP or irrelevant IgG antibodies as a negative control. Samples including the pre-enrich, EVs captured by the NW-PLAP and Dynabead-PLAP (bound), and the supernatants (unbound) collected after the immuno-enrichment were quantified for PLAP and CD63 proteins. The total measured PLAP proteins (sum of bound and unbound PLAP) in these samples were similar to that of the pre-enriched sample, indicating that there was minimal loss of PLAP (less than 12%) during the whole enrichment process (**Figure 5** A and B). PLAP+ve EVs were enriched with high efficiency by both the anti-PLAP conjugated NWs and Dynabeads, as 83.7±8.9% and 83.2±5.9% of the PLAP proteins were recovered respectively from the pre-enriched sample (Figure 5A and Figure S4A). These enrichments were significantly higher than the negative controls *(p* < 0.0001), which were 15.3±5% and 12.4±3.5% for the NWs and Dynabeads conjugated with an irrelevant antibody, respectively (Figure 5A). This data confirms that the enrichment of the PLAP+ve EVs with both NW-PLAP and Dynabead-PLAP is predominantly specific and mediated by the conjugated anti-PLAP antibodies. The PLAP+ve EV recovery performance can also be assessed using the ratio of bound PLAP protein (i.e. lysed from NW-PLAP / Dynabead-PLAP) relative to the total bound and unbound (supernatant) PLAP protein. Enrichments of 87.2±4.57% and 90.4±4.54% were obtained for anti-PLAP conjugated NWs and Dynabeads, respectively (Figure S4B). These recovery efficiencies are comparable to that reported by Menon *et al.* (95%) using the same clone of anti-PLAP (8B6) conjugated to protein-A agarose beads.^[51]^ It is noteworthy that in our study the irrelevant antibody controls captured non-specifically smaller amounts of PLAP proteins (17.1±5.27% and 14.1±2.3% for NWs and Dynabeads, respectively) (Figure S4B), compared to 30% reported by Menon *et al.* in plasma samples.

**Figure 5.**
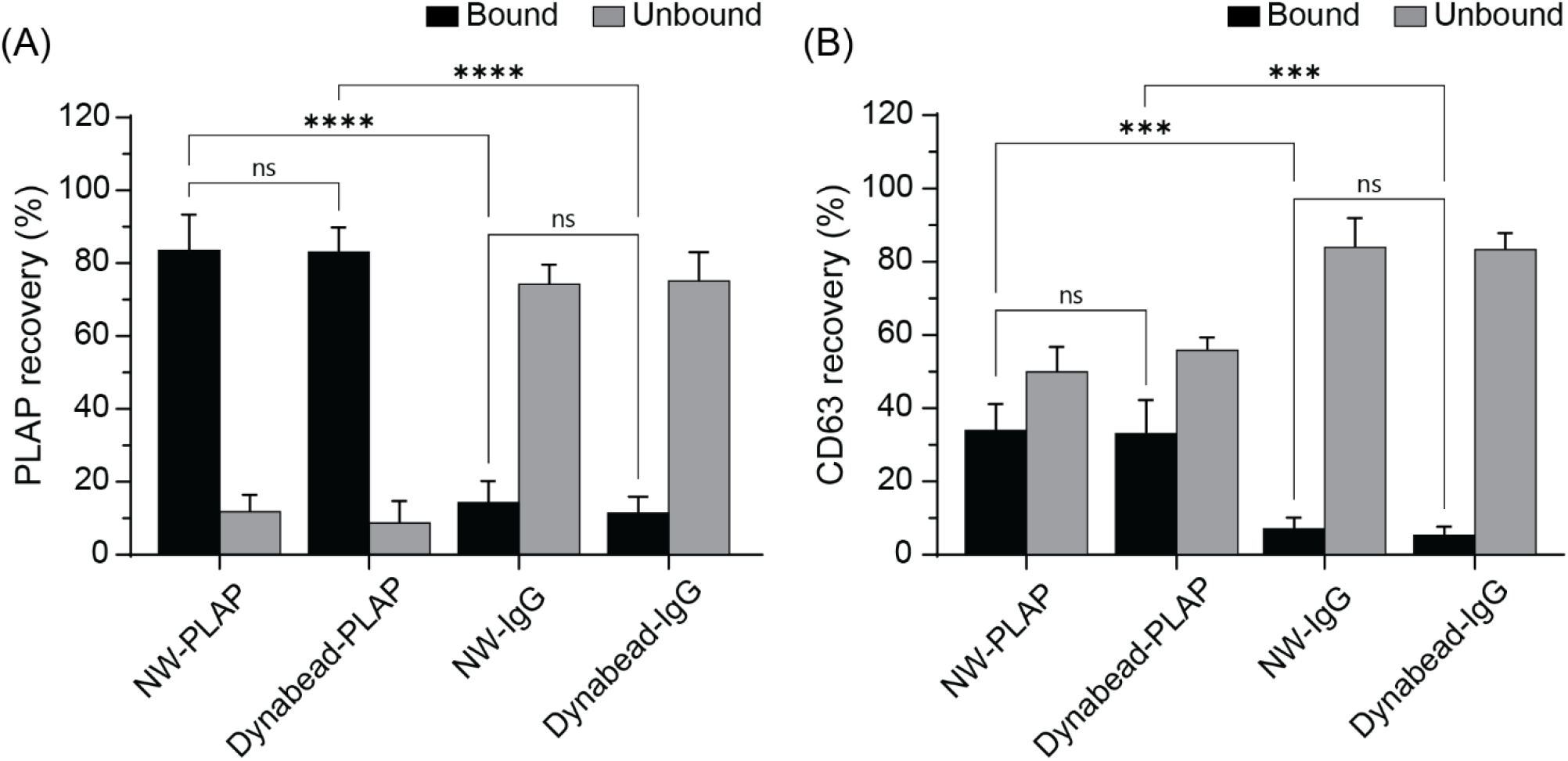
Recovery efficiency of (A) PLAP and (B) CD63 proteins extracted from EVs captured by NWs and Dynabeads conjugated with either anti-PLAP or an irrelevant IgG (bound) and the corresponding supernatants (unbound). Data are presented as means + SD *(n* = 3, one-way ANOVA with Tukey’s post-test, ns = not significant, \*\*\**p* < 0.001, and \*\*\*\**p* < 0.0001).

As a measure of the total EV population, we quantified the CD63 protein, a canonical marker expressed by most EV subtypes including both microvesicles and exosomes.^[47, 52]^ As approximately 30% of all EVs from BeWo cells are expected to express PLAP, CD63 recovery yields of 37.7% and 34.8% for the NW-PLAP and Dynabead-PLAP respectively, further confirmed successful enrichment of the subpopulation of PLAP+ve EVs (Figure 5B and Figure S4C). The NWs and Dynabeads conjugated with the irrelevant antibody controls yielded very low amounts of CD63 protein recovery with 7±3.2% for NWs and 5.4±2.2% for Dynabeads (Figure 5B), confirming that non-specific binding of EVs onto both NWs and Dynabeads is low. The significant reduction in the ratios of CD63 to PLAP proteins also observed for both NW-PLAP and Dynabead-PLAP from the pre-enriched sample further demonstrates that PLAP+ve EVs were specifically enriched with low contamination from non-target PLAP-ve EVs (Figure S4D). The purity of the target EV subtype (e.g. PLAP+ve EVs) and the purity of their protein/RNA molecular cargo is important to obtain reliable proteomic/transcriptomic data.^[53]^ Specifically, both target EV recovery yield and purity are important for insightful proteomic analyses with LC-MS.^[54]^

### 3.3 Proteomic analyses of PLAP+ve EVs enriched from BeWo cells

After demonstrating the potential of using iron oxide magnetic NWs to enrich PLAP+ve EVs, we further performed comparative proteomic analyses for the model EVs enriched from the culture media of BeWo cells by LC-MS/MS. As shown in Figure S5, ultracentrifugation followed by immuno-magnetic enrichment of EVs using the same weight of NW-PLAP and Dynabead-PLAP (1 mg) identified a total of 2822 and 2732 protein groups (in triplicate), respectively, 80.5% and 83.9% of which were enriched in all three replicates of NW-PLAP and Dynabead-PLAP, respectively. The number of proteins identified from EVs enriched from BeWo cell culture media was comparable to other previous reports using other approaches. For example, polyethylene glycol (PEG)-based exosome isolation from HeLa cell culture media identified 6299 proteins, 3254 (51.6%) of which were enriched in all three replicates.^[55]^

Using label-free quantification, we selected 2518 (89%) and 2545 (93%) protein groups that were identified in at least two out of three replicates for NW-PLAP and Dynabead-PLAP, respectively, for subsequent analyses. Cross-reference against the Vesiclepedia database showed that 95% of the selected protein sets could be identified as EV proteins (**Figure 6A**). As shown in Table S2, LC-MS/MS analyses also confirmed the presence of PLAP as well as several other markers commonly associated with EVs including transmembrane proteins (tetraspanins CD9, CD63, CD81) and several cytosolic membrane-associated proteins (TSG101, HSPA8, PDCD6IP, etc.). Pairwise intensity correlation of identified protein groups showed strong Pearson correlation coefficients for three replicates of either Dynabead-PLAP or NW-PLAP (0.987) as well as between their replicates (0.961), demonstrating not only the reproducibility of the EV enrichment methods but also the comparable enrichment performance of the NWs derived from biofilms of zetaproteobacteria *Mariprofundus ferrooxydans* to that of Dynabeads™, a commercial gold-standard (Figure 6B).

**Figure 6.**
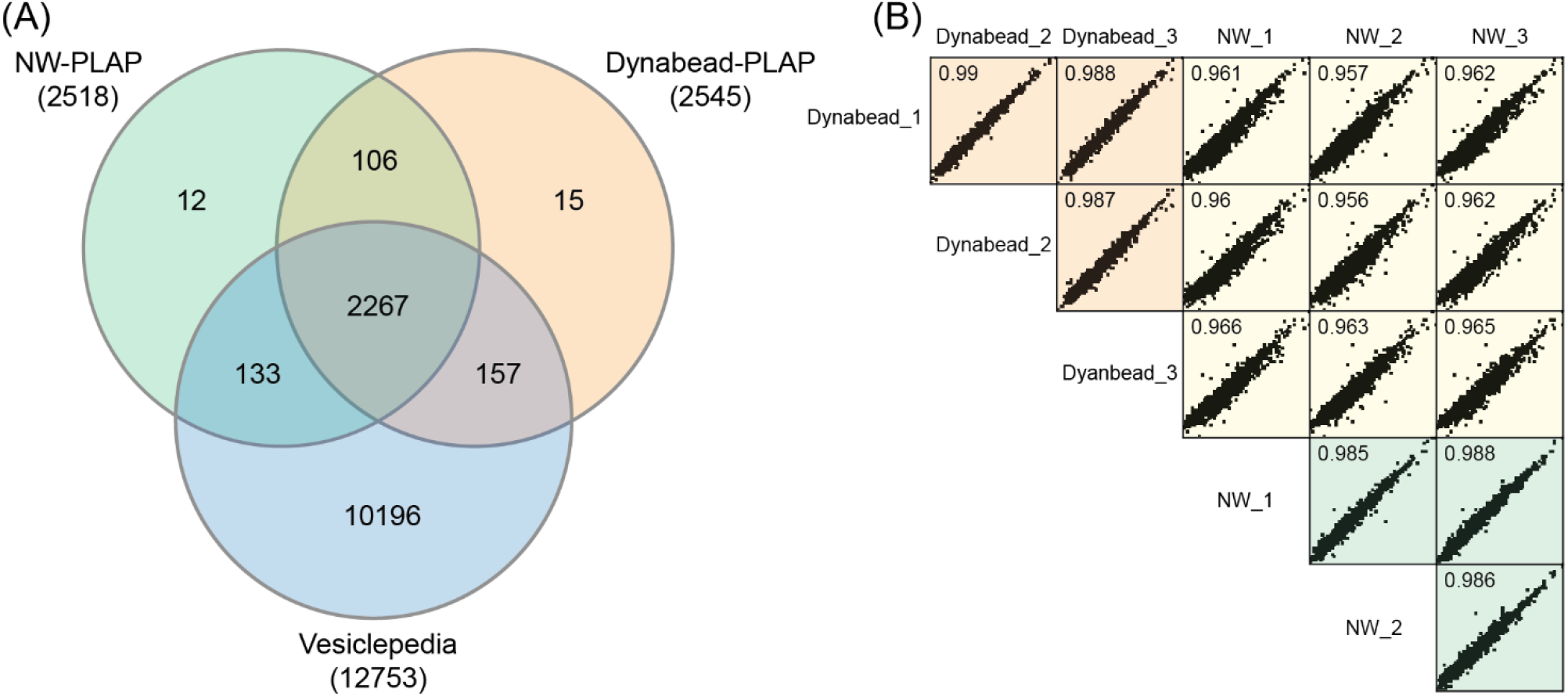
Comparative proteomic analyses of proteins identified in BeWo EVs enriched with NW-PLAP and Dynabead-PLAP. (A) Overlap of protein groups enriched with that of pre-enriched sample and Vesiclepedia database filtered to include EV proteins. (B) Pairwise correlation of intensities of protein groups enriched within replicates of NW-PLAP and Dynabead-PLAP as well as between their replicates. Inset numbers indicate Pearson correlations.

Detailed information about the cellular components of enriched proteins was further acquired by analyzing gene ontology (GO, DAVID Bioinformatics Resources).^[56]^ As shown in Figure S6A, GO analysis demonstrated that the extracellular exosome category was the most highly enriched in the NW-PLAP sample, while this category ranked second for samples enriched with Dynabead-PLAP after the cytosol category. Specifically, among the identified protein groups enriched by Dynabead-PLAP and NW-PLAP, 36% and 37.8% were respectively found to be located in the extracellular exosome category (GO:0070062) while the majority of proteins were categorized into the cytoplasm (GO:0005737), cytosol (GO:0005829), and nucleus (GO:0005634) categories (Figure S6B). This result is attributed to the fact that PLAP proteins were demonstrated to be detected in microvesicles (100–1000 nm) and exosomes (50–200 nm) isolated by ultracentrifugation at 10,000 × *g* and 100,000 × *g*, respectively.^[57–58]^ Microvesicles have been reported to carry several proteins from the cytoplasm and nucleus.^[59]^ The insights obtained from GO analyses agreed with those obtained from the Vesiclepedia database and NTA result, demonstrating the ability of the magnetic NWs to specifically enrich PLAP+ve EVs.

## 3. Conclusion

We report an immuno-magnetic enrichment approach for specific EV subtypes and their subsequent processing for high quality mass spectrometry. Specifically, we have demonstrated comparable EV-enrichment yield and purity using biofilm-derived iron oxide NWs to that of Dynabeads™, using PLAP+ve EVs as a model for a physiologically relevant EV subtype. In addition, low non-specific binding of non-targeted EVs (7% for NW) was achieved. The captured EVs could be readily eluted or directly lysed for downstream analyses. We subsequently optimized an EV mass spectrometry method after immunomagnetic enrichment with both NWs and Dynabeads™, which consistently yielded a high number of EV proteins. The immuno-specific EVs enrichment for mass spectrometry approach using naturally occurring iron oxide magnetic NWs or Dynabeads™ provides an efficient method for EV proteomic studies.

## 4. Experimental Section

### Materials and Reagents

1-Ethyl-3-(3-dimethylaminopropyl)carbodiimide hydrochloride (EDC-HCl), N-hydroxysuccinimide (NHS), 4-morpholineethanesulfonic acid monohydrate (MES monohydrate), ethanolamine, bovine serum albumin (BSA) were purchased from Sigma Aldrich. Anti-PLAP antibody (8B6) was purchased from Life Technologies Pty Ltd.

### Preparation of Iron Oxide NWs

Bacteria-derived NWs were extracted and purified as described previously.^[38, 46]^ Briefly, bacteria biofilm waste provided by SA Water (South Australia) was resuspended in milli-Q water and purified through repeated washing and centrifuge cycles to remove organic and inorganic impurities. The purified NWs extracted from the biofilms were annealed at 600 °C to change their crystal phase and increase their magnetic properties. After thermal annealing, the NWs were washed again with milli-Q water, dried at 70 °C and stored at room temperature until use.

### Preparation of Poly(methacryloyloxyethyl phosphate) 3-block-poly(acrylamide)_20_ RAFT Polymer (RAFT-3MAPC2-20AAm)

RAFT-3MAPC2-20AAm diblock polymers, referred to as *p*AAm, were synthesised based on a previously published procedure with small modifications.^[60]^ Briefly, the RAFT diblock polymer consisted of RAFT-COOH agent (2-(((butylthio)carbonothioyl)-thio)-propanoic acid), anchoring (methacryloyloxy)ethyl phosphonate groups (n = 3 monomers), and a block of stabilising polyacrylamide polymer (n = 20 monomers).

### Conjugation of Antibody to NWs and Dynabeads

The NWs were first coated with the *p*AAm diblock polymers. NWs (10 mg) and *p*AAm (50 mg) were first resuspended in milli-Q water and sonicated. The pH of *p*AAm solution was adjusted to 4 using 0.1 M sodium hydroxide and added dropwise to the NW suspension while being sonicated on ice. The mixture was left to react under continuous mixing (300 rpm) on an orbital shaker for 18 h at room temperature. After that, the pH of the mixture was sequentially adjusted to pH 5 and then 6 using 0.1 M sodium hydroxide with icecold sonication in between. *p*AAm-coated NWs (NW-*p*AAm) were washed with milli-Q water three times to remove salts and unreacted copolymers and finally resuspended in milli-Q water to the desired concentration of 10 mg mL^-1^. Carbodiimide coupling reaction was carried out to covalently conjugate antibodies to the NW-*p*AAm (Scheme 1A) through the terminal carboxylic acid functional groups. Briefly, NW-*p*AAm (1 mg), EDC-HCl (76 mg mL^-1^) and NHS (11.5 mg mL^-1^) were mixed in milli-Q water (mixture pH 5.7) for 5 min at room temperature. The activated NWs were then subsequently washed once with milli-Q water and phosphate buffer saline (1 × PBS pH 7.4, 10010023, Life Technologies) and immediately added to PBS buffer containing the antibodies (40 μg). The conjugation reaction was carried out under continuous mixing (800 rpm) for 2 h at room temperature using a ThermoMixer (Eppendorf). The resulting antibody-conjugated NWs were washed three times with PBS buffer to remove non-reacted antibodies, further reacted with 50 mM ethanolamine to quench non-reacted activated carboxylic acid groups and blocked with 0.5% BSA. The antibody-conjugated NWs were finally resuspended in PBS buffer, stored at 4 °C and used within two weeks of conjugation. The conjugation efficiency of antibodies to NWs was determined by:

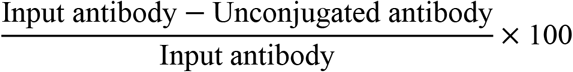

Dynabeads M-270 carboxylic acid (14305D, Invitrogen) were used as the reference gold standard. The beads were activated and conjugated to antibodies according to the manual. Briefly, Dynabeads (3 mg) were washed with 25 mM MES buffer and reacted with EDC-HCl (50 mg mL^-1^) and NHS (50 mg mL^-1^) with shaking for 30 min at room temperature. Activated beads were subsequently washed with 25 mM MES buffer and PBS buffer and reacted with antibodies (120 μg) for 2 h at room temperature. Antibody-conjugated Dynabeads were washed twice with PBS buffer, quenched with 50 mM ethanolamine, and blocked with 0.5% BSA. The beads were resuspended in PBS buffer, stored at 4 °C and used within two weeks of conjugation.

### Characterization of Iron Oxide NWs

The morphology of the NWs was analysed by field emission scanning electron microscope (FE-SEM, Carl Zeiss Microscopy Merlin) and TEM (Tecnai G2 operated at 120 kV and equipped with an EDAX X-ray detector). The relative evolution of the NW sizes upon biofunctionalization was monitored by measuring changes in the scattering profile with DLS (Zetasizer Nano ZS90, Malvern). The elemental composition was measured using XPS (Kratos AXIS Ultra DLD) for NWs casted onto silicon wafer substrates as described previously.^[61]^ XPS survey scans (0–1200 eV) were performed with a pass energy of 160 eV and a step size of 0.5 eV. High-resolution spectra of C 1s, O 1s and N 1s were scanned with the pass energy of 20 eV and step size of 0.1 eV. XPS spectra were analyzed using the CasaXPS software (v2.3.18). The binding energy of the survey and high-resolution scan was referenced to that of C 1s CC/CH peak at 284.8 eV. The magnetic property of these iron oxide NWs derived from biofilms was evaluated by a vibrating sample magnetometer (VSM) using a magnetic field of ± 20 kOe.

### Cell Culture and Pre-enrichment of Total EVs

BeWo choriocarcinoma cells (trophoblast-like cell line) were cultured at 37 °C in a humidified incubator with 5% CO_2_ in Roswell Park Memorial Institute media (RPMI 1640, Gibco) supplemented with 10% fetal bovine serum (FBS, Gibco) and penicillin/streptomycin (10 000 U mL^-1^, Gibco). Once 80–90% confluency was reached, the culture media was removed, and the cells were washed twice with 1 × Dulbecco’s phosphate-buffered saline (DPBS, Gibco). The cells were then continuously cultured in the FBS-free medium for an additional 48 h. Culture media was collected and total EVs were pre-enriched based on a previously published protocol.^[62]^ First, the media containing EVs (22 mL) was centrifuged at 300 × *g* for 20 min and 2000 × *g* for 20 min at 4 °C (5699-R rotor, Rotanta 460 R, Hettich) to remove dead cells and debris. The supernatant was then centrifuged at 10 000 × *g* for 30 min at 4 °C to pellet large EVs and then ultracentrifuged at 120 000 × *g* for 70 min at 4 °C (70 Ti rotor, Optima L-80 XP, Beckman Coulter) to pellet small EVs. Pellets of small and large EVs were washed twice with PBS buffer, combined together, resuspended in PBS buffer and stored at −80 °C for two weeks.

### Enrichment of PLAP+ve EVs by NWs and Dynabeads

Pre-enriched BeWo EVs were incubated with comparable amounts of anti-PLAP antibody conjugated magnetic NWs or Dynabeads (NW-PLAP or Dynabead-PLAP) overnight at 4 °C with shaking (800 rpm) to promote attachment of EVs. EV-captured NWs and Dynabeads were separated using 0.5 T neodymium magnets (3489, Aussie Magnet) fixed into a 3D-printed magnetic separation rack (Figure S7), and the supernatants were collected for the quantification of PLAP proteins. EV-captured NWs and Dynabeads were washed three times with 500 μL of PBS buffer. EVs were either released from the magnetic NWs/Dynabeads for nanoparticle tracking analysis (NTA) or lysed directly to extract the protein cargo (Scheme 1B). After applying the magnet, the lysates were collected and stored at −80 °C for ELISA and mass spectrometry analysis.

### Nanoparticle Tracking Analysis

The concentration and size of EVs were characterized by NTA (Nanosight NS300, Malvern). EVs enriched by NWs and Dynabeads were eluted using 0.1 M Glycine-HCl pH 2.8 with an incubation of 30 min at room temperature. After magnetic removal of NWs and Dynabeads, the supernatant containing eluted EVs was collected and neutralized by the addition of 1 M Tris pH 8. EVs were diluted in ultrapure water to obtain approximately 50 particles per frame for optimal counting. All measurements were performed in triplicate and each sample was analyzed five times with the same camera settings (shutter speed: 30.15 ms, camera level: 15, capture duration: 60 s).

### Quantification of PLAP and CD63 Proteins from Enriched EVs

After incubating pre-enriched EVs with NW-PLAP or Dynabead-PLAP, the captured EVs were lysed with 1 × RIPA buffer (Merck Millipore, 0.5 M Tris-HCl pH 7.4, 1.5 M NaCl, 2.5% deoxycholic acid, 10% NP-40, 10 mM EDTA) containing 1 × cOmplete™ Mini EDTA-free Protease Inhibitor Cocktail (Roche) for 20 min at 4 °C. As a control, pre-enriched EVs were directly lysed without isolation. Lysates were quantified for PLAP and CD63 proteins using a human PLAP ELISA kit (MBS289869, MyBiosource) and a human CD63 ELISA kit (EH95RBX5, Thermo Fisher Scientific). The recovery of PLAP protein was calculated by dividing the amount of PLAP or CD63 proteins extracted by that of the pre-enriched sample.

### Proteomic Analysis

Sodium dodecyl sulfate polyacrylamide gel electrophoresis (SDS-PAGE) of extracted EV proteins followed by in-gel digestion of proteins were used to process the samples for mass spectrometry analyses. EV-captured NWs and Dynabeads were resuspended in 1 × RIPA buffer containing 1× protease inhibitor as described above, and the lysates were collected after removing the NWs and Dynabeads with a magnet. Prior to in-gel digestion, the total protein concentration was quantified by bicinchoninic acid (BCA) assay (Pierce™ BCA Protein Assay Kit, Thermo Fisher Scientific). EV protein extracts were mixed with 4X NuPAGE™ LDS sample buffer (Invitrogen) and heated at 70 °C for 10 min prior to running on NuPAGE™ 4–12% Bis-Tris gels (Invitrogen) for 5 min to ensure the complete removal of remnant NWs and Dynabeads. Areas of gel containing proteins were then excised in reference to a prestained protein ladder, with those areas subsequently cut into smaller pieces prior to performing an in-gel digestion method adapted from the gel-aided sample preparation (GASP) protocol.^[63]^ Briefly, gel pieces were dehydrated with acetonitrile and dried at room temperature. Protein-containing gel pieces were then reduced with 10 mM dithiothreitol (Sigma) in 100 mM ammonium bicarbonate at 56 °C for 45 min, dehydrated again with acetonitrile, and then alkylated with 55 mM 2-chloroacetamide (Sigma) in 100 mM ammonium bicarbonate in the dark at room temperature for 30 min. After removing the liquid, gel pieces were rehydrated with 5 mM ammonium bicarbonate and dehydrated again in acetonitrile prior to re-hydration with enzymatic digestion (Trypsin/Lys-C mix, Promega) in 5 mM ammonium bicarbonate overnight at 37 °C. Peptides were sequentially extracted from the gel using 1% formic acid (40 μL), 50% acetonitrile in 1% formic acid (50 μL), and acetonitrile alone (100 μL) followed by sonication of 15 min in between before supernatants were collected. Peptide-containing extracts were vacuum-dried and resuspended in 0.1% formic acid prior to liquid chromatography-tandem mass spectrometry (LC-MS/MS) analysis.

LC-MS/MS analyses were conducted on an EASY-nLC 1200 system (Thermo Scientific) coupled to an Orbitrap Exploris 480 mass spectrometer (Thermo Scientific). Five microlitres of the peptide sample were loaded onto a 50 °C heated C18 fused silica column (75 μm × 25 cm) at a flow rate of 600 nL min^-1^. The column was packed with 1.9 μm ReproSil-Pur 120 C18-AQ particles (Dr. Maisch) and possessed a pulled tip. Peptide chromatography was performed over a 70 min linear gradient (3–20% acetonitrile in 0.1% formic acid) with a flow rate of 300 nL min^-1^. Two compensation voltages (–50 and –70 V) were alternately applied from a FAIMS Pro interface (Thermo Scientific) to regulate the entry of ionized peptides into the mass spectrometer. MS scans *(m/z* 300 to 1500) were acquired at resolution 60000 *(m/z* 200) in positive ion mode (2 kV spray voltage). MS/MS scans were measured at resolution 15000 after multiply charged peptide precursors (minimum intensity 10000) were sequentially fragmented with higher energy collision dissociation (HCD) at 27.5% normalized collision energy. A dynamic exclusion period of 40 s was specified, and cycle times were restricted to 1.5 s.

The raw LC-MS/MS data files were subjected to label-free quantification (LFQ) using Maxquant (v1.6.2.6) with peptide spectra searched against the human UniProt database (downloaded on 4/2021) using the integrated Andromeda search engine.^[64]^ For the identification of peptides, the digestion enzyme was set to Trypsin/P with a maximum of two missed cleavages, a fixed modification of cysteine carbamidomethylation, and variable modifications of protein N-terminal acetylation and methionine oxidation. Mass tolerances of 20 ppm and 0.1 Da were used for precursor ions and fragment ions, respectively. The false discovery rate (FDR) was 1% for both peptides and proteins. LFQ data were processed with Perseus (v1.6.2.1) to determine correlations between the sample triplicates. Protein groups identified by LC-MS/MS were compared with the Vesiclepedia database and Functional Enrichment analysis tool (FunRich v3.1.3). A Venn diagram was generated by InteractiVenn tool, and gene ontology (GO) analyses were performed using the database for annotation, visualization and integrated discovery (DAVID, v6.8).^[56]^

## Supporting Information

Supporting Information is available online or from the author.

## Acknowledgements

The authors would like to acknowledge the financial support of the Thrasher Research Fund (Award Number: 15144) and the National Health and Medical Research Council Development Grant (APP1171821). B.T. was supported by National Health and Medical Research Council Investigator Grant (GNT1197173). D.L. and T.T. acknowledge the support from the Australian Research Council, ARC Research Hub for Graphene Enabled Industry Transformation, funding under Industrial Transformation Research Hub (grant IH150100003). The authors also acknowledge Bioplatforms Australia, and the State and Federal Governments, which co-fund the NCRIS-enabled Mass Spectrometry and Proteomics Facility at the University of South Australia.

---

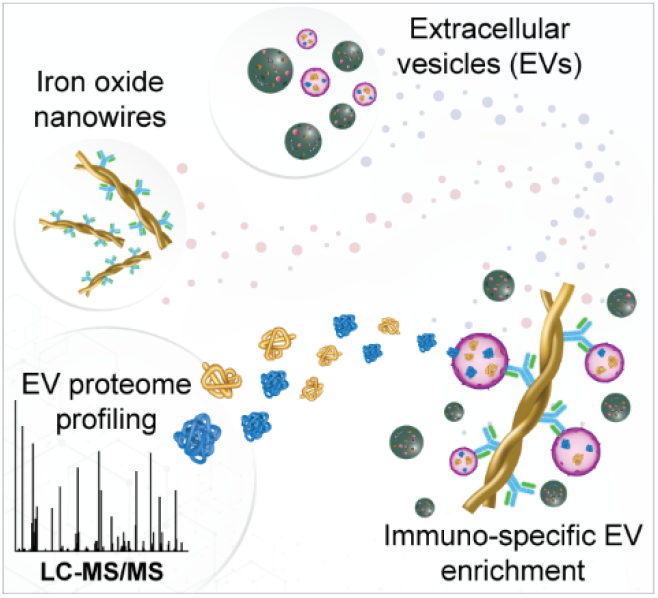

A magnetic enrichment method of immuno-specific extracellular vesicles (EVs) for mass spectrometry-based proteome profiling was developed using iron oxide nanowires (NWs) extracted from bacteria biofilms. By covalently conjugating NWs with antibodies, specific EV subtypes were efficiently enriched with low contaminations of proteins and non-specific bindings of non-target EVs, which enables reproducible quality EV analysis using liquid chromatography-tandem mass spectrometry.

