## Supporting information for "Magnetic Enrichment of Immuno-Specific Extracellular Vesicles for Mass Spectrometry Using Biofilm-Derived Iron Oxide Nanowires" (from another version of this preprint: version 1, 2022-05-01)

C. Young, P. Hoffmann

Clinical and Health Sciences, University of South Australia, City West Campus, Adelaide, South Australia 5000, Australia

N. T. H. Pham

School of Chemistry, The University of Sydney, Sydney, New South Wales 2006, Australia

T. T. Tung, D. Losic

School of Chemical Engineering and Advanced Materials, The University of Adelaide, Adelaide, South Australia 5005, Australia

* Denotes co-corresponding author

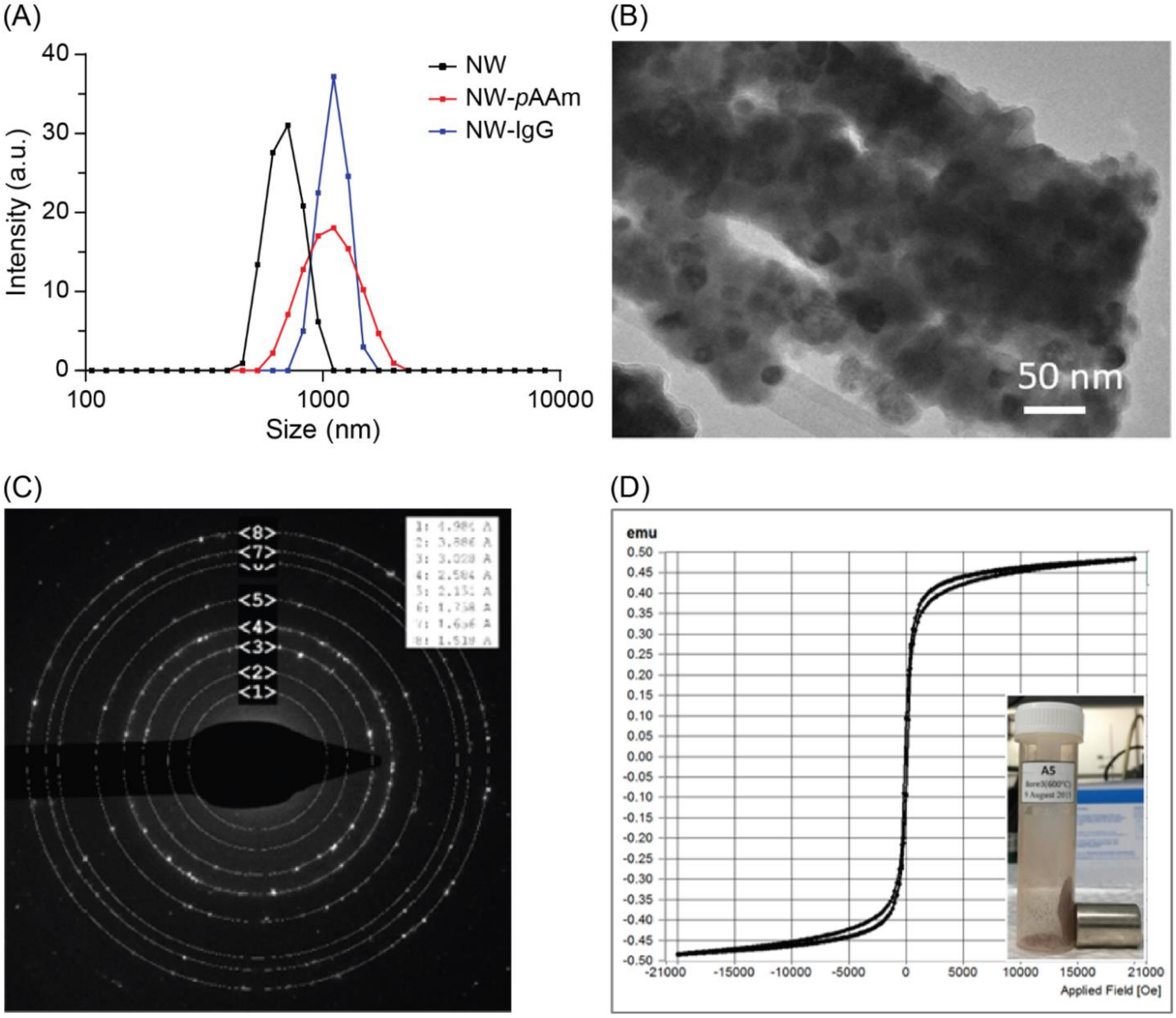


**Figure S1.** (A) Monitoring of changes in NWs' scattering by dynamic light scattering during the coating procedure. (B) Transmission electron microscope image and (C) diffraction pattern of iron oxide NWs showing wire-like structures with diameters of 50–80 nm and polycrystalline rings with lattice refractions corresponding to a mixture of magnetite and hematite structures. (D) The magnetic hysteresis curves of iron oxide NWs revealing a ferromagnetic behaviour of the material with a photo (inset) demonstrating the magnetic property of iron oxide NWs using a magnet.

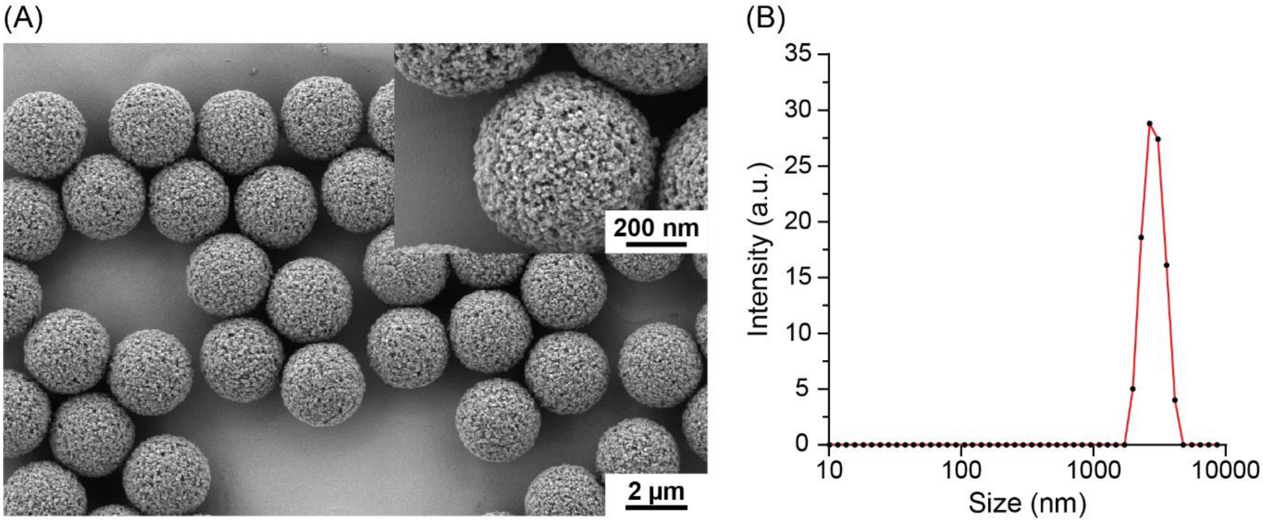


**Figure S2.** (A) Scanning electron microscopy images of Dynabeads M-270 carboxylic acid. Inset is the SEM image at a higher magnification. (B) Dynamic light scattering size measurement of Dynabeads™.

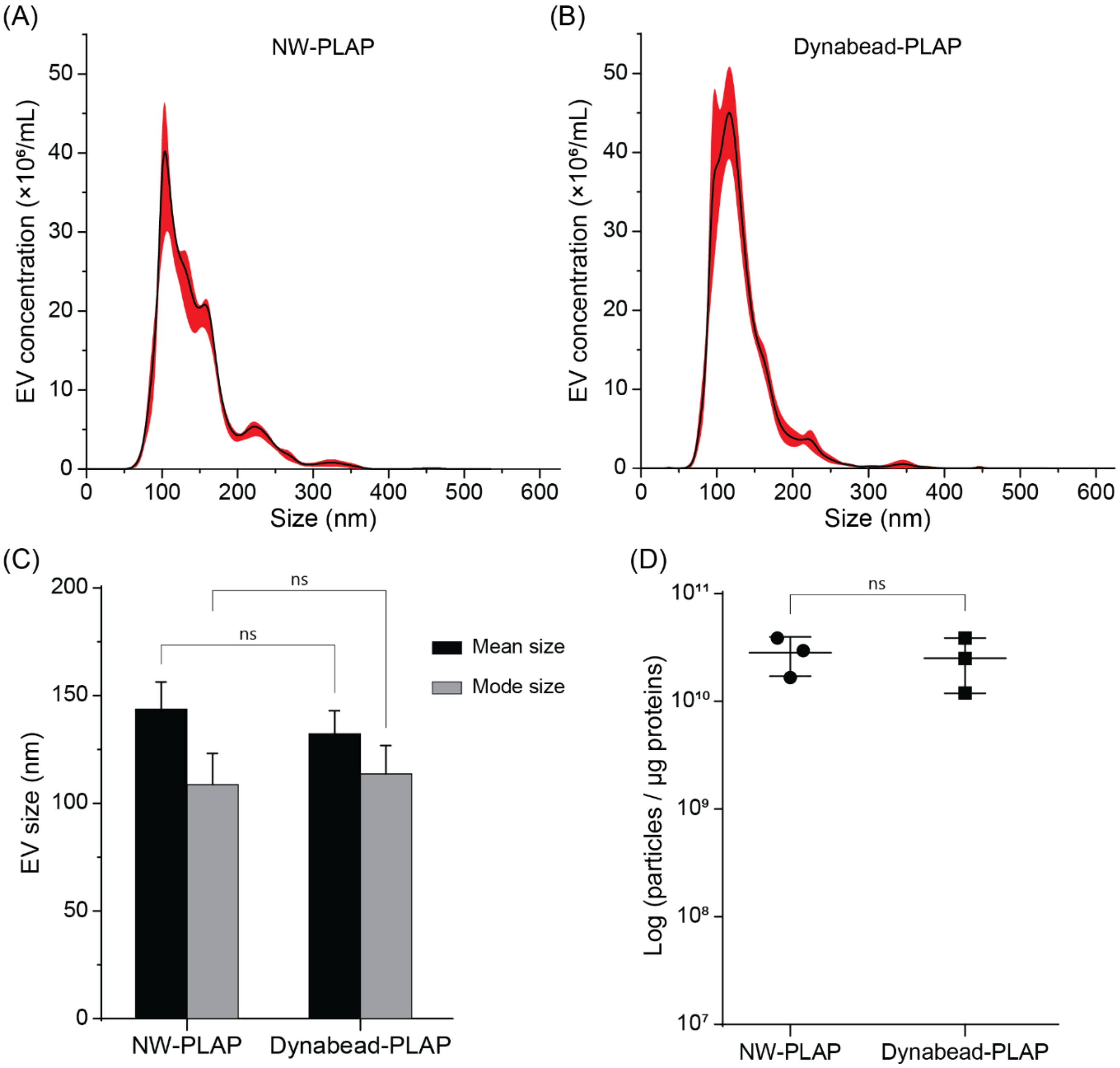


**Figure S3.** Nanoparticle tracking analysis of eluted EVs enriched by (A) NW-PLAP and (B) Dynabead-PLAP. (C) Mean size and mode size of eluted EVs. (D) The particle to protein ratio of PLAP+ve EVs enriched by NW-PLAP and Dynabead-PLAP. Data are presented as means + SD (n = 3, independent Student's t-test with 95% confidence interval, ns = not significant, $p > 0.05$).

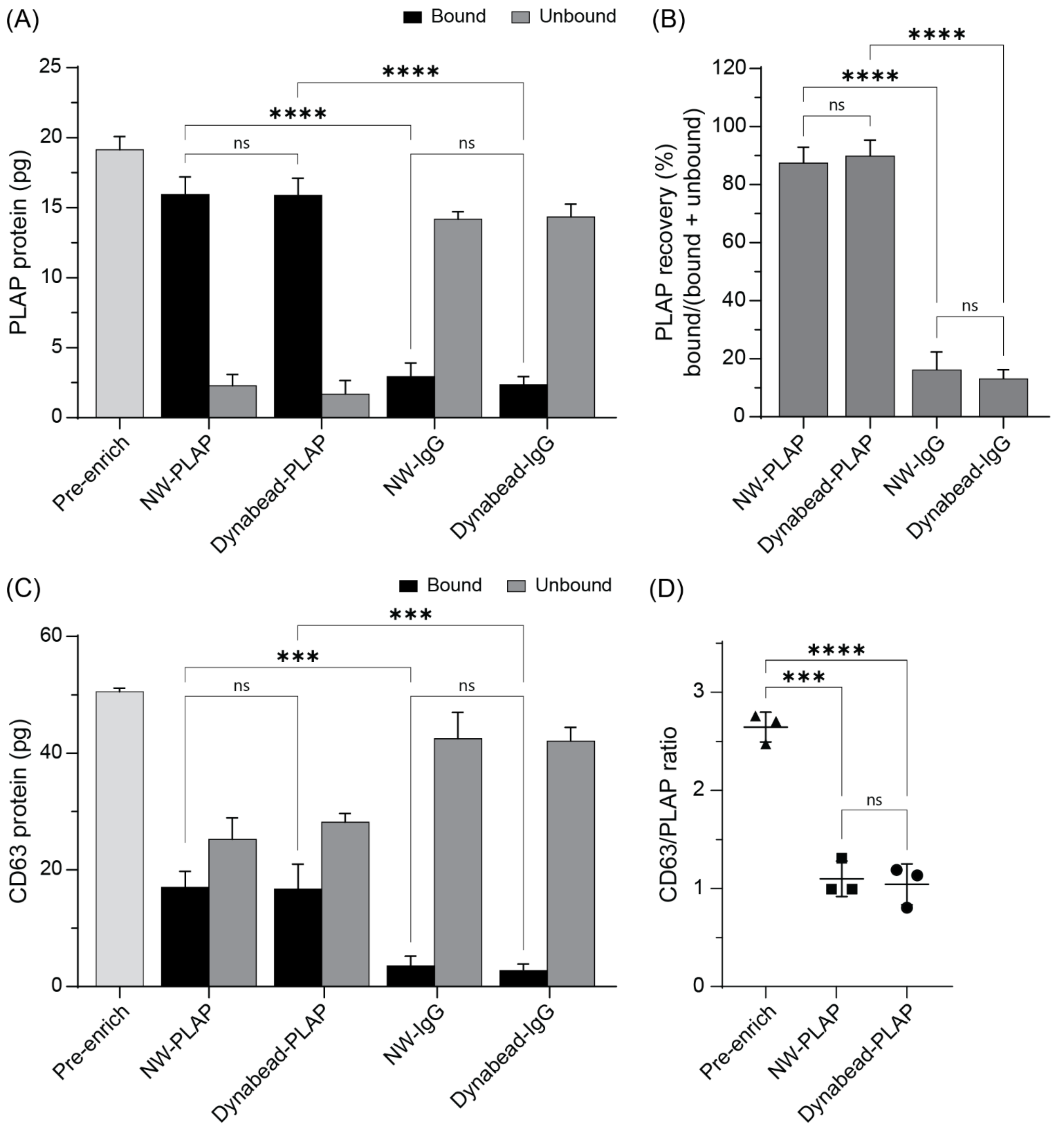


**Figure S4.** (A) Quantification of PLAP in the pre-enriched sample (after ultracentrifugation of BeWo cell culture media), EVs captured by NWs and Dynabeads conjugated with either anti-PLAP or irrelevant IgG (bound) and the corresponding supernatants (unbound). (B) PLAP recovery of EVs enriched by NW-PLAP and Dynabead-PLAP presented by the ratio of bound to the total of bound and unbound. (A) Quantification of CD63 in the pre-enriched sample, bound and unbound of NW-PLAP and Dynabead-PLAP. (D) The ratio of CD63 to PLAP (pg/pg) calculated for the pre-enriched sample, NW-PLAP and Dynabead-PLAP. Data are presented as means + SD (n = 3, one-way ANOVA with Tukey's post-test, ns = not significant, ***$p < 0.001$, and ****$p < 0.0001$).

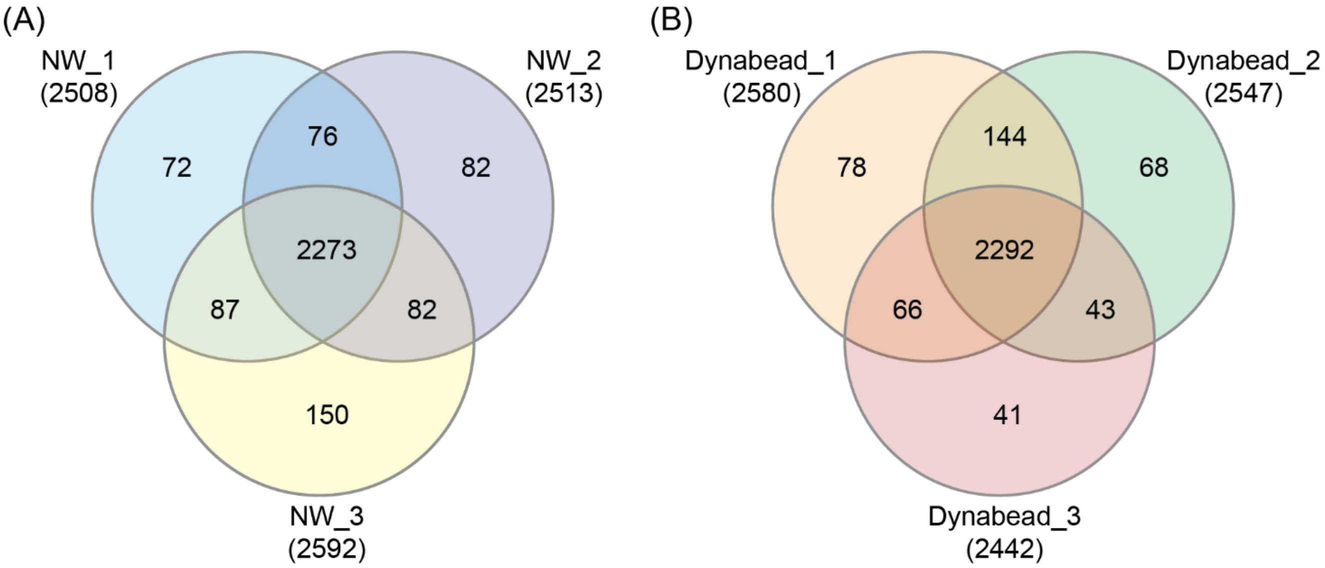


**Figure S5.** Overlap of protein groups enriched by three replicates of (A) NW-PLAP and (B) Dynabead-PLAP.

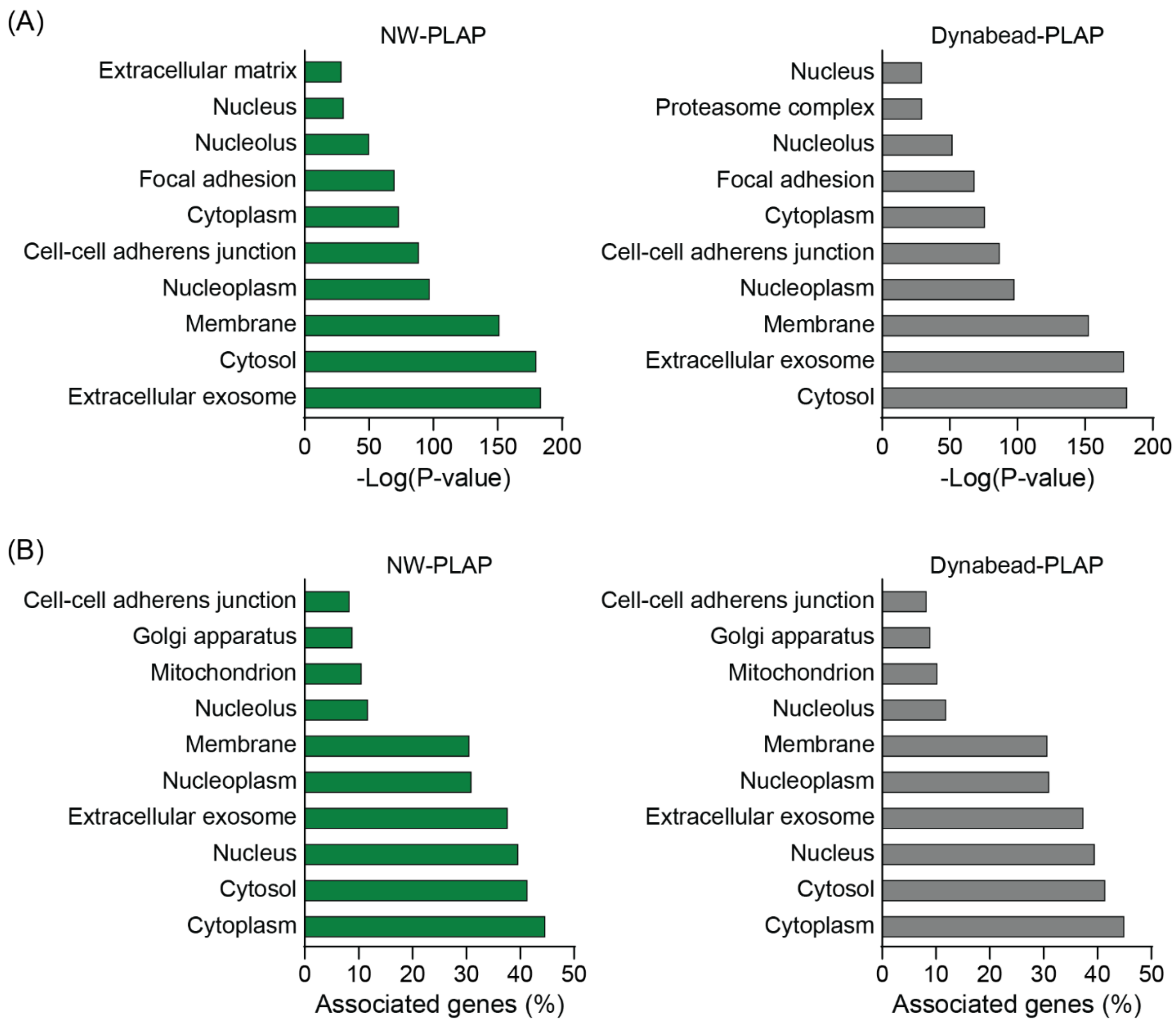


**Figure S6.** Gene ontology (GO) analyses of the protein groups in BeWo EVs enriched by NW-PLAP and Dynabead-PLAP according to DAVID Bioinformatics Resources. Top 10 GO categories were reported for (A) Cellular component and (B) The percentage of associated genes.

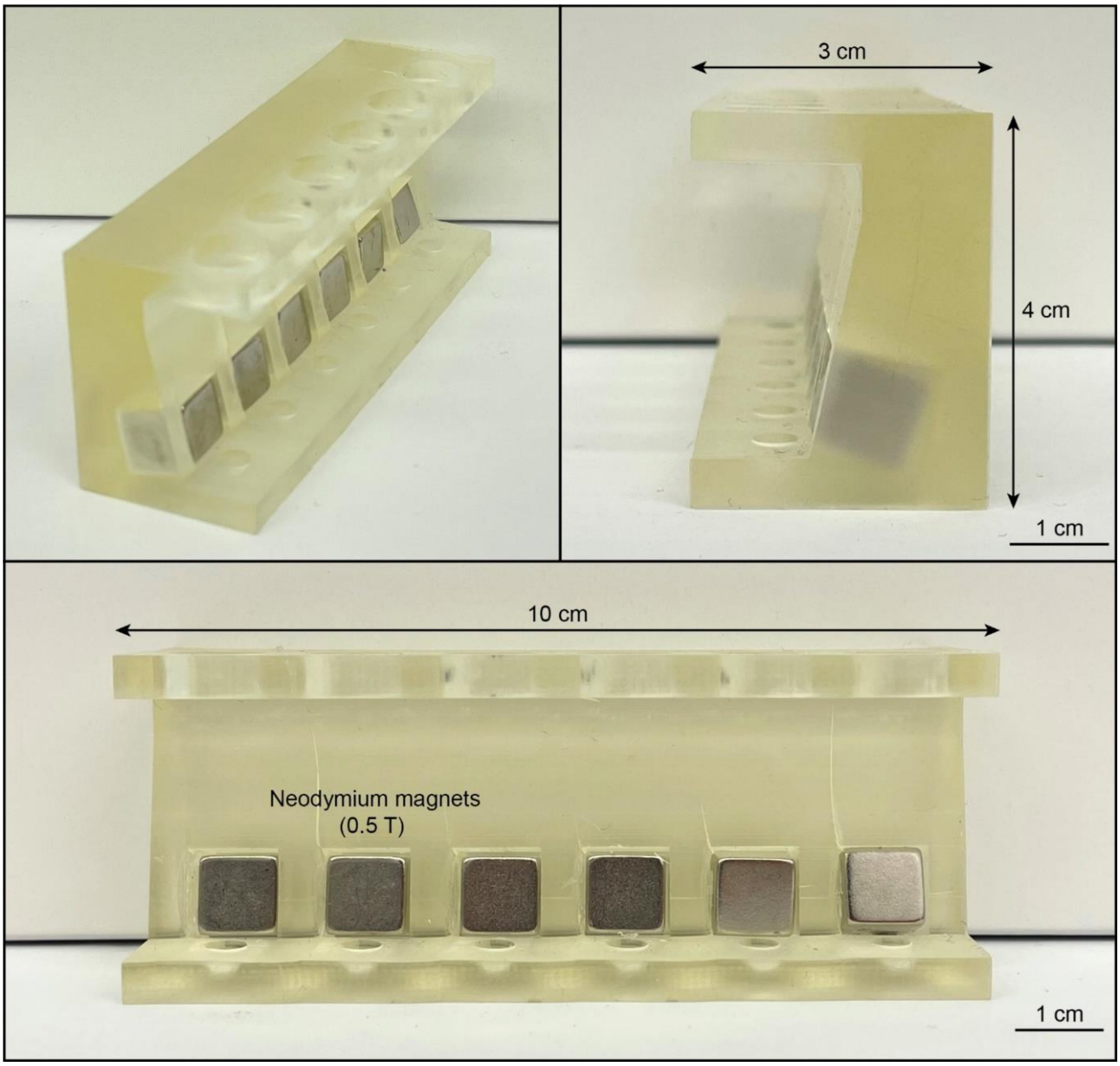


**Figure S7.** Images of the in-house 3D-printed magnetic separation rack used for immuno-magnetic isolation of EVs. The rack was designed to fit 1.5-mL and 2.0-mL Eppendorf tubes and neodymium magnets (Grade N42, 0.5 T, 10 × 10 × 10 mm).

**Table S1.** Batch-to-batch validation of antibody conjugation to NWs

| NW samples | XPS elemental composition (%) | | |
|---|---|---|---|
| | C | N | O |
| NW Batch 1<br>Antibody conjugation 1 | 47.79 | 8.13 | 44.09 |
| NW Batch 1<br>Antibody conjugation 2 | 45.80 | 7.51 | 46.69 |
| NW Batch 2<br>Antibody conjugation 1 | 49.12 | 7.85 | 43.03 |
